# Competition between plasmids conferring the same antimicrobial resistance leads to plasmid-host network specialisation

**DOI:** 10.1101/2025.05.09.652947

**Authors:** Arthur Newbury, Angus Buckling, Elze Hesse, Dirk Sanders

## Abstract

Plasmids have a key role in disseminating antimicrobial resistance (AMR). Previous work has shown that in the presence of antibiotics, plasmids conferring AMR will spread to a larger number and a more phylogenetically diverse group of host organisms. However, this pattern may be complicated when multiple plasmids confer the same resistance.

Here, we analyse host-plasmid networks in experimental communities of three bacterial species cultured for six weeks with three incompatible plasmids (plasmids that cannot stably coexist within a single cell) conferring tetracycline resistance. The communities are cultured either without tetracycline, or with a low or high dose of the antibiotic. We find that while antibiotic selection leads to higher plasmid prevalence, it does not change the structure of host-plasmid networks. Under all conditions, the networks become highly specialised, as specific host-plasmid pairings emerge.

This result is supported by a previously published mathematical model parameterised to match our experimental set-up. Our study reiterates the importance of fitness effects of plasmids in determining the structure of bacteria-plasmid networks. The spread of a given AMR plasmid is reduced when fitness differences are reduced due to other plasmids carrying AMR genes, leading to more specialised bacteria-plasmid networks.

## Introduction

Predicting and mitigating the spread of Antimicrobial resistance (AMR) is a key public health challenge, as antibiotic-resistant infections will have devastating costs on human society as both debilitating and lethal diseases increase (Naghavi et al., 2024; O’Neill, 2016)The main vectors responsible for the dissemination of AMR genes are plasmids, common mobile genetic elements that can transfer between unrelated bacterial hosts (Castañeda-Barba et al., 2024; Dimitriu, 2022). Antibiotics can make plasmids that carry AMR genes highly beneficial for bacteria, with consequences for their prevalence and spread (Carroll & Wong, 2018) even if some bacterial host populations subsequently evolve AMR when plasmid encoded AMR genes are transferred to the host chromosome (MacLean et al., 2025). Bacteria and plasmids form complex interaction networks as plasmids of the same type can be hosted by multiple bacterial strains, and individual bacterial populations (and indeed individual bacteria) may host multiple distinct plasmids (Klümper et al., 2015; Lee & Ko, 2021; Madec & Haenni, 2018; Stalder et al., 2019)). Thus, predictions and methods from network ecology can be used to characterise and predict trends in bacteria-plasmid transmission dynamics. Specifically, bacteria-plasmid interactions can be represented with bipartite networks which consider only the links between (but not within) two types of interactants – here plasmids and bacteria. These approaches have been instrumental in recent work highlighting the effects of antibiotics on both experimental and natural bacteria-plasmid networks(Newbury et al., 2022; Risely et al., 2024).

One key network metric is connectance - the proportion possible interactions that are actually realized. Highly connected networks are in general associated with fast transmission dynamics. In bacteria-plasmid networks the spread of genes may be exacerbated by a number of super spreaders (Shapiro et al., 2023). Generality of a species within a network measures not only the number of interactions but how evenly distributed they are amongst other species. Thus, generalist species populations are less sensitive to large changes in the population sizes of one or a few other species (Kehoe et al., 2021). A link between AMR carrying plasmids and high connectance in bacteria-plasmid systems has been demonstrated in a complex natural microbial community (Risely et al., 2024). The study shows that in wastewater beneficial broad-host range plasmids carrying AMR genes are associated with a wider range of hosts. Furthermore, that generality and connectance emerge *because* of a mutualistic host-plasmid relationship (plasmid relying on bacteria for replication and bacteria relying on plasmid for AMR) has been experimentally demonstrated in simplified experimental communities with only a single plasmid (Newbury et al., 2022). This suggests that AMR plasmids are important drivers of gene exchange between bacteria and plasmids.

However, this pattern may be counteracted when multiple plasmids confer the same resistance. There are two reasons to suspect this, illustrated by a simplified situation with a single bacterial strain and two types of plasmid that confer resistance to the same antibiotic that is present in the local environment. 1 In intraspecific competition between cells hosting either plasmid, the fitness differences are reduced (relative to the case where only one plasmid confers resistance). 2 A cell that contains one type of plasmid may gain little or no fitness benefit from receiving the second type of plasmid (though possibly at very high antibiotic concentrations, having more copies of the resistance gene will be beneficial (Dimitriu et al., 2021; Wellner et al., 2025)). These two subtly different and complementary processes will play out in tandem, meaning that each plasmid is no longer almost guaranteed to persist at high prevalence or at all within the host population (or spread through several hosts populations if they were available) as would be expected if it was the only one conferring resistance. However, process 1 is potentially the more important of the two, since the reduction of fitness differences between competing strains does not require that both plasmids confer resistance to the *same* antibiotic. Here, we isolate and explicitly investigate process 1 by using plasmids that cannot co-exist within the same cell.

Plasmids are classified into incompatibility (Inc) groups (Garcillán-Barcia et al., 2023) where the plasmids in each Inc group share elements of their partition or replication systems that prevent stable coexistence within a single cell (Novick, 1987). For example, the majority of pB plasmids extracted by Droege et al.(Dröge et al., 2000) from wastewater belong to the Inc-P group and confer tetracycline resistance. In these circumstances, AMR plasmids that would be expected to become prevalent within individual host populations and spread throughout the microbial community may instead be outcompeted by other beneficial plasmids, potentially weakening the relationship between plasmids conferring AMR and both generality and connectance in bacteria-plasmid networks. Though experimental evidence is lacking on this point.

Here we investigate how the presence of AMR plasmids with overlapping resistance profiles affects the formation of host-plasmid network structures under different antibiotic selection pressures. We test this with experimental microbial communities containing three bacterial species (isolated from compost *Bordetella sp*., *Ochrobactrum sp*. and *Pseudomonas sp*.) and the broad host range, Inc-P plasmids pB5, pB12 and pKJK5::GFP. We then use a host plasmid population dynamics model to generalise our results beyond the specific bacteria and plasmids used in the experiment.

## Results

All three plasmids confer resistance to the antibiotic tetracycline and become beneficial to their hosts in the presence of low (1 μg/m) or high (1 μg/m) concentrations of tetracycline (Figure 1). As a result, tetracycline at either concentration had a large effect on the total plasmid prevalence within the community (Figure 2a). While there was a decrease in plasmid prevalence from week 1 to 6 in the No Tet treatment, plasmids were still detected in all replicates after 6 weeks. However, there were marked differences between hosts: *Pseudomonas sp*. was largely fully infected with pB12 in week 6 regardless of the treatment, whereas *Ochrobactrum sp*., which tended to host pB5, maintained a low plasmid prevalence in the absence of tetracycline (Figure 3c) but typically carried plasmids when tetracycline was present. *Bordetella sp*. showed considerable variation between replicates (Figure 2b) and across treatments. However, due to its lower number (i.e. number of *Bordetella* sp. colonies per agar plate), these estimates are expected to be more varied. Thus, the differences in plasmid prevalence between treatments and across time result largely from differences in plasmid prevalence within *Ochrobactrum sp*. populations, which dominated in all treatments (Figure 3).

**Figure 1.**
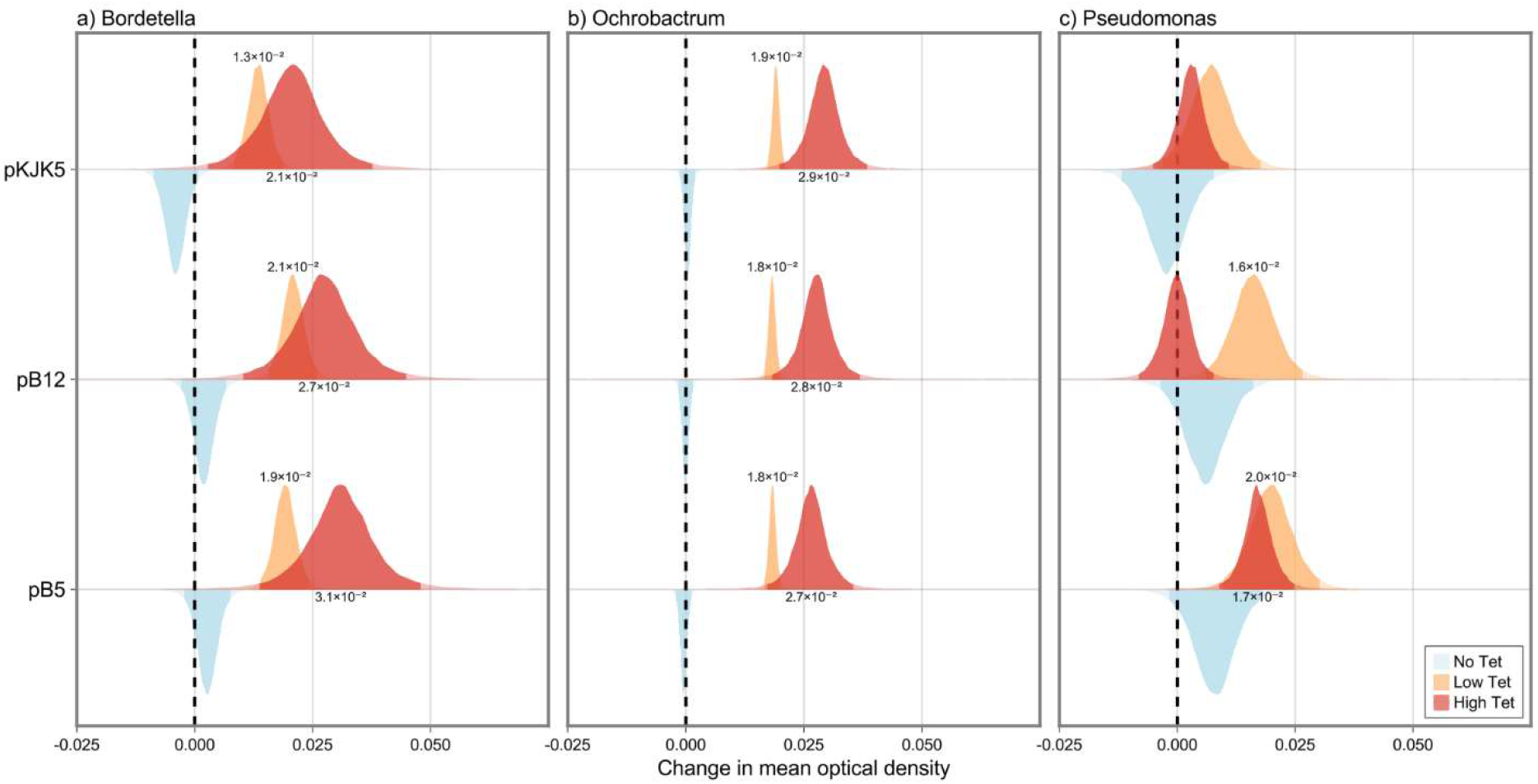
Estimated effect of three plasmids on individual fitness of three bacterial hosts in the 3 treatments: No Tet, Low Tet and High Tet, based on optical density readings. Since the 3 bacterial strains took different lengths of time to reach carrying capacity, area under the curve (AUC) of the logistic growth function fit to optical density data were divided by the total time of the growth assay for the given strain, giving units of mean optical density, comparable between strains. Shown are posterior estimates of the effect each plasmid had on the respective host strains’ mean optical density. All significant effects (95% credible intervals not containing 0) labeled with point estimate (posterior median) with labels for Low Tet and High Tet above and below the density plots respectively. Mineral oil was used avoid evaporation during growth assays. Under these more anerobic conditions, the High Tet treatment was able to inhibit the growth of *Pseudomonas sp*. populations even when hosting a plasmid. Despite this, *Pseudomonas sp*. was able to survive in the main experiment in the Hight Tet treatment.

**Figure 2.**
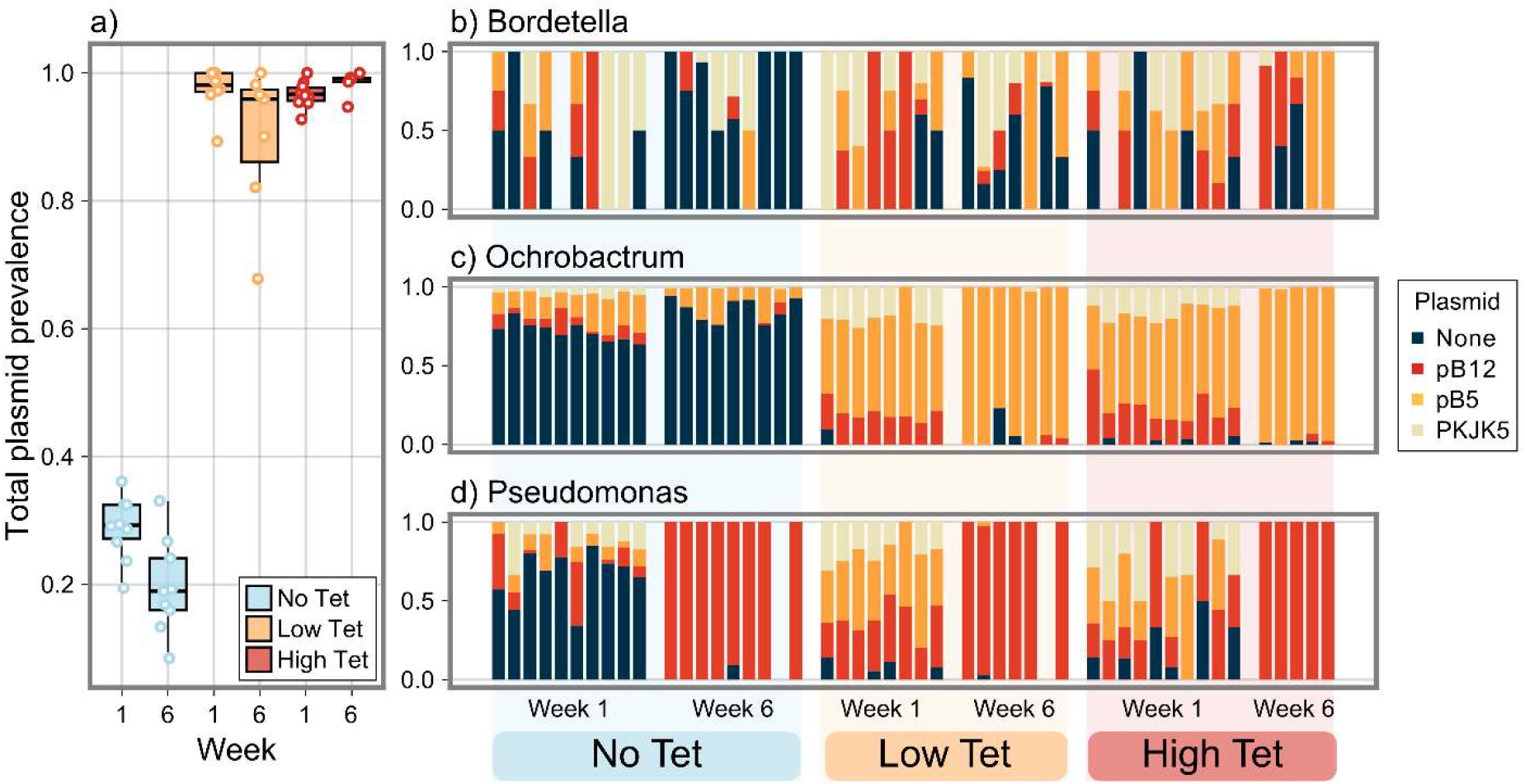
Plasmid prevalence. a) Overall plasmid prevalence in weeks 1 and 6 for treatments with no, low and high tetracycline concentration plotted are individual values, the median and interquartile range. b-d) Proportions of each plasmid per host per microcosm.

**Figure 3.**
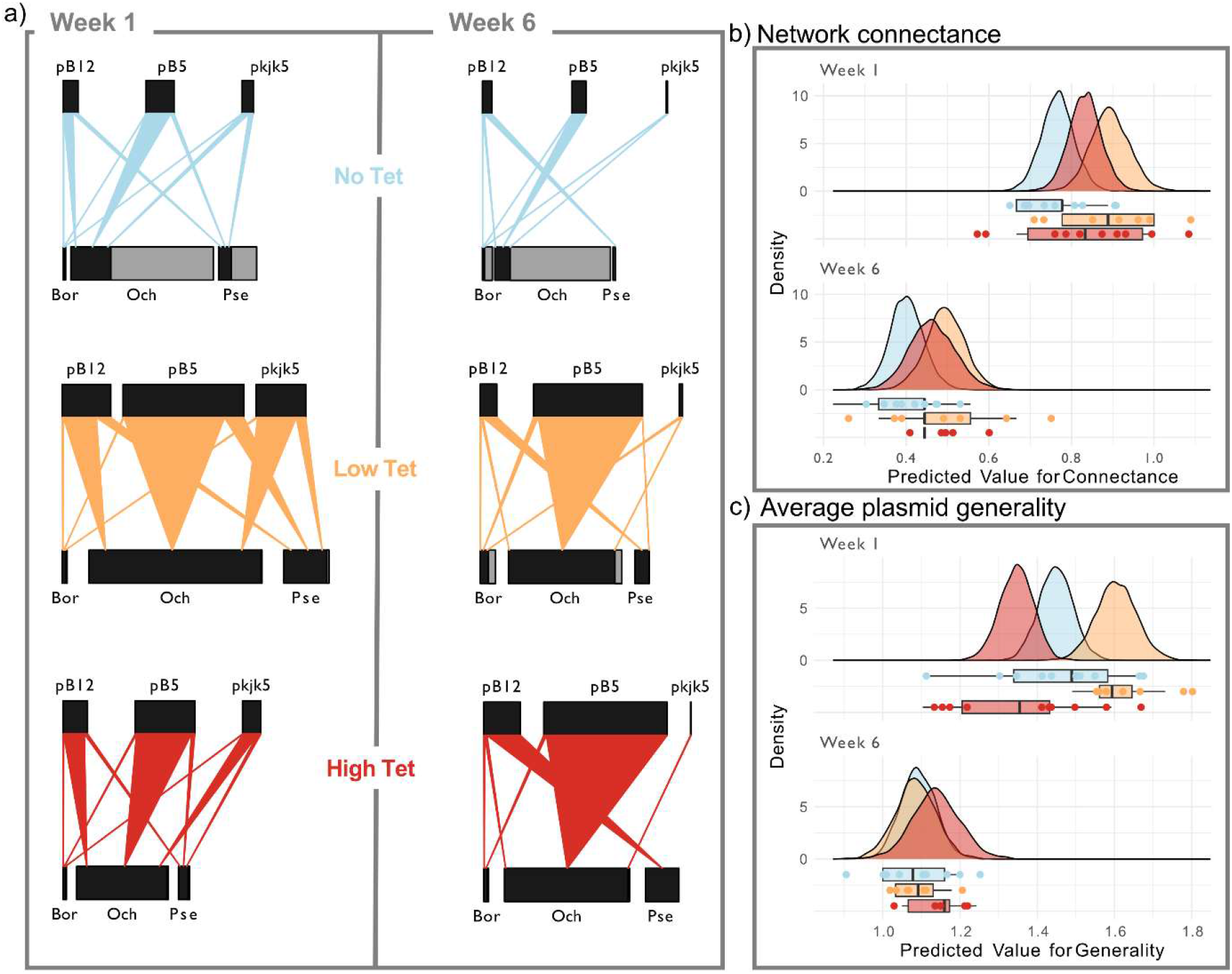
Experimental host-plasmid networks. a) Bipartite networks with bacterial hosts as lower and plasmids as higher network level in weeks 1 and 6 for the treatments with no, low and high tetracycline concentration. Plotted are treatment means for relative bacterial abundance and infection rate per host-plasmid combination. The size of the lower bar represents relative bacteria abundance in relation to the other bacteria, with the grey part indicating the proportion of each bacterial population that is not associated with any plasmid. Network connectance b) and average plasmid generality (across all three plasmids) c) in weeks 1 and 6 for the three different treatments, plotted are the posterior distributions from bmrs models testing for the interactive effect of treatment and week on each variable. Below the axis are individual data points, the median and interquartile range for the observed values for each treatment from the experiment.

Theory and empirical data suggest that beneficial plasmid networks (here, the Low Tet and High Tet treatments) will exhibit higher connectance (number of realized host-plasmid associations divided by the number of possible associations) and generality (the extent to which plasmids interact with multiple species to a similar extent). Therefore, we analysed the response of these network metrics to the treatments across time. Starting from fully and evenly connected networks (see Methods), host-plasmid networks in all treatments became highly specialized by the end of the experiment (Figure 3a). Both connectance and average plasmid generality declined significantly between weeks 1 and 6 (Figure 3bc). Interestingly, though generality dropped to similar levels across all treatments it dropped off faster in the High Tet treatment (Figure 3c). This was similar for all three plasmids (Figure 4a-c), however, pB12 appeared to stay more general (Figure 4a). Connectance tended to be lowest in the No Tet treatment for both weeks 1 and 6 (Figure 3b). However, there were no statistically significant differences between treatments *within* a given time point (see overlap in posterior distributions). Thus, the final networks were not significantly different in terms of the generality and connectance. In particular, average plasmid generality (i.e. generality across all three plasmids), which is more robust to total prevalence of plasmids during sampling than connectance, was remarkably similar across treatments (Figure 3c).

**Figure 4.**
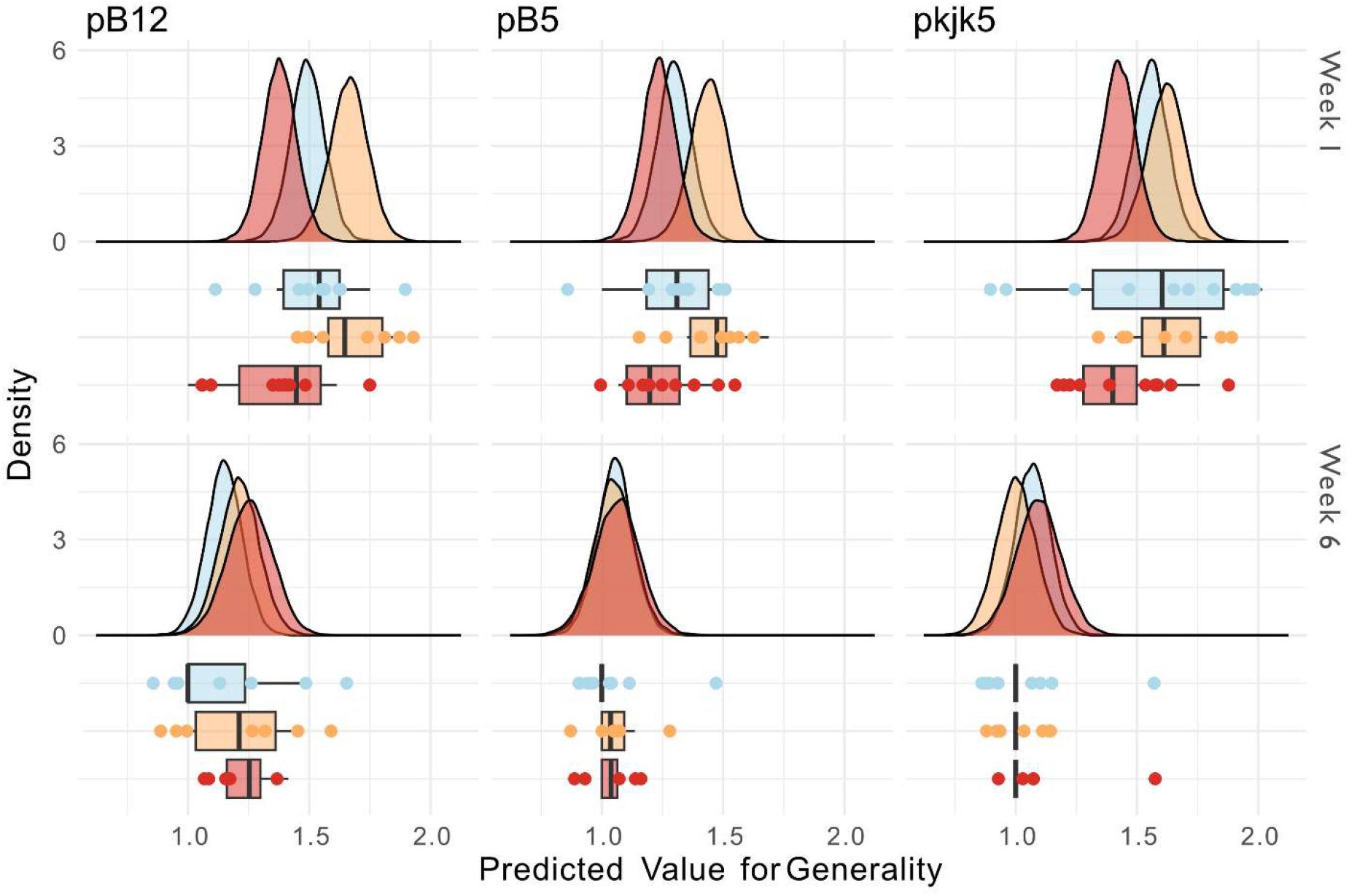
Generality for the plasmids (a) pB12, (b) pB5 and (c) pkjk5 in weeks 1 and 6 for the three different treatments (no, low and high tetracycline concentration), plotted are the posterior distributions from bmrs models testing for the interactive effect of treatment and week. Below the axis are individual data points, the median and interquartile range for the observed values for each treatment from the experiment.

Plasmid presence impacted host community composition, just as host community dynamics had a large impact on the prevalence of specific plasmids. Alongside the plasmid containing communities, we also cultured identical bacterial communities without any plasmids, under the same conditions (No Tet, LowTet, High Tet). For both the plasmid and no plasmid communities we used Dirichlet-multinomial models to determine the proportions of each host, plasmid for each treatment and time point, while accounting for differing total cell densities between replicates (Figure 5). *Ochrobactrum sp*. was the dominant host in weeks 1 and 6 in all treatments apart from High Tet without plasmids, where *Bordetella sp*. was the most prevalent strain. As a result, pB5 was also the most successful plasmid across all treatments due to being the plasmid most commonly found with *Ochorobactrum sp*..

**Figure 5.**
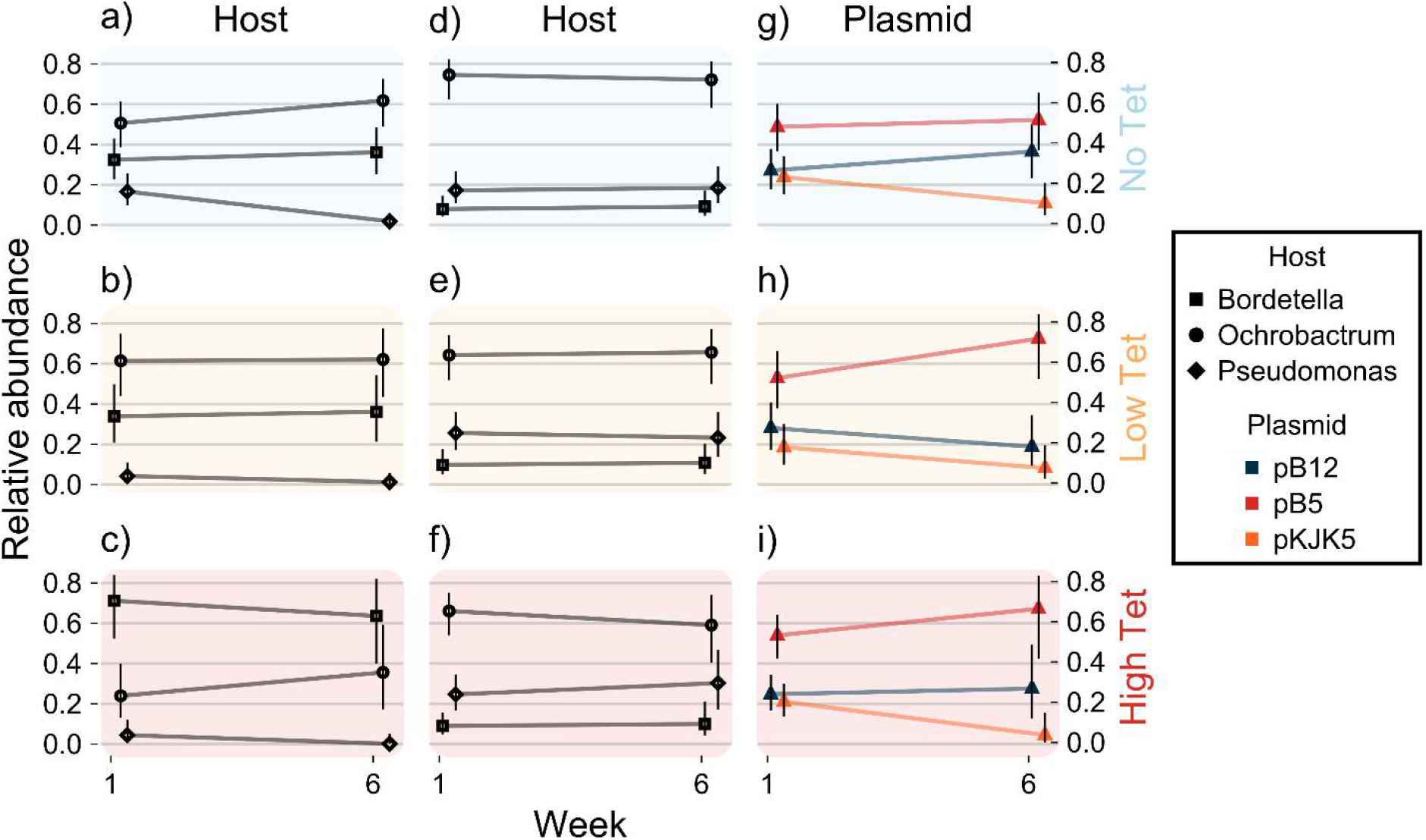
Community dynamics. Relative abundance of bacterial hosts, plasmids and their host-plasmid combinations in the three antibiotic treatments in week 1 and 6. a-c) Relative abundance of hosts without plasmids present. d-f) Relative abundance of hosts with plasmids present. g-i) Relative abundance of plasmids. Points denote posterior medians from a Dirichlet-multinomial model based on colony counts, with error bars indicating 95% credible intervals.

The main result - that the networks became highly specialised regardless of the antibiotic treatment - is at odds with earlier theoretical and empirical work suggesting that positive fitness effects (here tetracycline resistance in the Low Tet and High Tet treatments) should lead to greater generality and connectance. Therefore, we parameterised a plasmid population dynamics model from (Risely et al. 2024) to approximate our experimental set up - multiple incompatible plasmids, all or none conferring a benefit to all hosts. Specifically, we ran simulations with 3 hosts and 3 plasmids, where each plasmid had a negative impact on the growth rate of each host, i.e., all plasmids were costly. Then we ran alternative simulations with all parameters kept the same except that the growth rate effects of the plasmids were uniformly increased such that now all plasmids increased the growth rates of all hosts, i.e., all plasmids were beneficial. By contrasting the results between these simulations, we were able to determine the impact of beneficial plasmids on network structure when all plasmids are beneficial. We then compared these results with those where we repeated the same process as above, but only one plasmid was made beneficial. As quantities in the model may approach but never actually reach 0, we use a cutoff of 1% infection rate to determine whether or not a plasmid interacts significantly with a host and calculate a plasmid’s connectance as the number of these significant interactions divided by 3.

Unlike earlier results for the case where one or a subset of plasmids is *beneficial* to a subset of hosts (Figure 6a), costly plasmids that interacted with few hosts (Figure 6b) tended to still interact with few hosts after growth rate effects were modified to make all plasmids beneficial (Figure 6c). As a result, costly plasmids with a connectance of 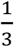 (only interacting with a single host population) had a 96.5% chance of having the same connectance when they were beneficial (Figure 6d). By contrast, when only a single plasmid was made beneficial it almost always had a connectance of 1 (Figure 6e). This indicates that the tendency for plasmids to interact with more hosts when they confer fitness benefits is diminished by competition with other plasmids that also benefit the host bacteria.

**Figure 6.**
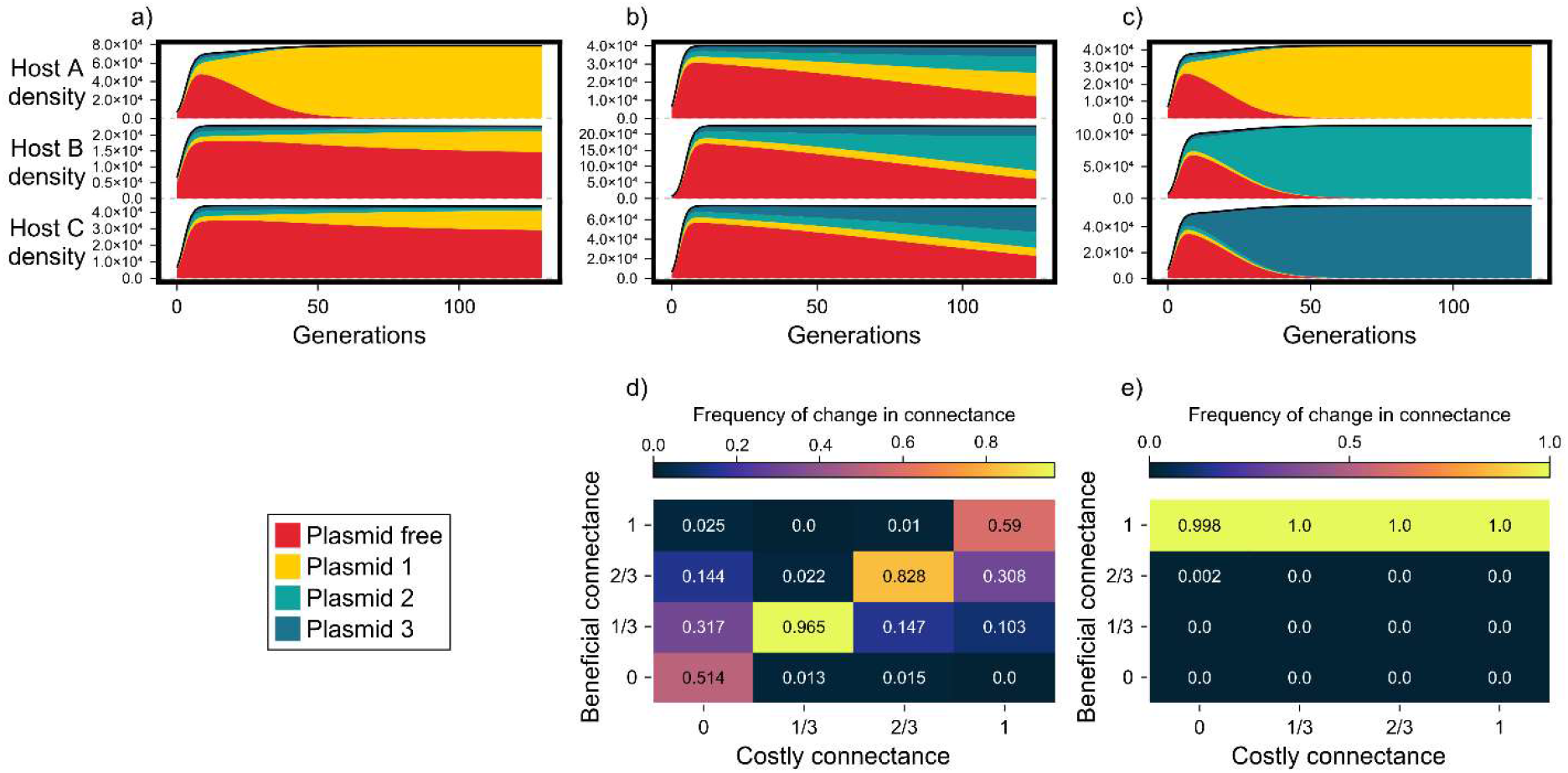
Community dynamics model. a-c) depict examples of host plasmid dynamics. a) Plasmid 1 benefits Host A, while all other interactions are costly. Here, a single beneficial interaction leads to generality of the beneficial plasmid, even though it only benefits a single host. b) All plasmids are costly. Hosts specialise on the plasmid least costly to them. c) All plasmids are beneficial; hosts specialise on the plasmid that is most beneficial to them. If the least costly plasmid for a given host is also the most beneficial in another context (e.g. the presence of antibiotics) then the change in context will not affect network structure. d-e) show how a plasmid’s connectance changes when it goes from being costly to beneficial. The value in each cell is the frequency with which plasmids with a given connectance (costly connectance) changed to the corresponding connectance when they became beneficial (beneficial connectance). d) When all plasmids become beneficial to all hosts, the connectance of any given plasmid is most likely to remain the same. This is most pronounced for specialised plasmids (connectance = ⅓). Otherwise, when a plasmid’s connectance does change, there is typically a shift towards intermediate values. e) When only a single plasmid becomes beneficial to all hosts it will almost always have a connectance of 1.

## Discussion

The host-plasmid networks in our experimental communities became less connected and plasmids more specialist over time, with specific host-plasmid pairs emerging by week 6. The emerging specialised networks were remarkably similar across treatments with and without antibiotics, despite plasmids being selected for and reaching high prevalence only in presence of antibiotcs. This contrasts with results from previous theoretical models and experimental results showing that the generality of a single plasmid is driven by presence and absence of the antibiotic the plasmid confers resistance to and therefore the plasmid either being beneficial or costly (Newbury et al., 2022). Our modelling suggest our results can be simply explained by the decreased variance in fitness effects of plasmids in both the presence and absence of antibiotics. Furthermore, the results highlight that prevalence and generality in plasmids are not necessarily linked, even with increased prevalence the plasmids in this experiment became more specialised..

The networks that emerged by the end of the experiment were all highly specialised, with particularly prevalent host-plasmid combinations being stable in both presence and absence of antibiotics. Thus, plasmid prevalence was determined by individual host-plasmid interactions (Figure 3) and not by the level of plasmid generality. For example pB5 being the most prevalent overall due to its high prevalence within *Ochrobactrum* populations, which was in turn the most prevalent host across all plasmid-containing treatments. In fact, in the absence of tetracycline, pB5 only infected a small portion of *Ochorobactrum sp*., but was still more prevalent than pB12 which typically fully infected *Pseudomonas sp*.. Despite being highly prevalent within the dominant host under antibiotic selection, pB5 did not spill over and become prevalent within the other host populations as we would have expected in the absence of other tetracycline resistance plasmids. (This prediction is drawn from previous experimental and theoretical work (Hurtado et al., 2024; Newbury et al., 2022; Risely et al., 2024).

There was interspecific variation in terms of generalism as well as in the specific host-plasmid combinations, though this may partly be due to differences in host cell densities. *Bordetella sp*. stayed by far the most generalist host, but due to its generally low abundance this had little effect on the overall structure of the networks. *Ochrobactrum sp*. was already predominantly hosting the plasmid pB5 by week 1, while *Pseudomonas sp*. was far more generalist at week 1. This may be due to differences in growth rate - *Ochrobactrum sp* had reached higher cell densities by week 1, meaning more opportunity for deterministic patterns to emerge. Unlike the other two hosts, *Pseudomonas sp*. was typically fully infected (with pB12) even without tetracycline by week 6.

This last in particular highlights In line with previous work (Newbury et al., 2022) and the model presented here, we would expect different networks to emerge if only one of the 3 plasmids conferred tetracycline resistance. Also, if the plasmids came from different incompatibility groups, we may have seen increased generality in all treatments due to reduced competition between plasmids. However, further experimentation is needed to confirm this. Mathematical modelling has played an important role in understanding plasmid ecology and making broad predictions (Alonso-del Valle et al., 2021; Newbury et al., 2022; Stewart & Levin, 1977). Given the complexities of plasmid biology such as post-segregational killing systems, plasticity in transmission rates, rapid evolution and systems preventing host susceptibility to other plasmids (Dimitriu, 2022), it is not possible to confidently make more focused predictions (e.g. the sub-network that will emerge from a small number of closely related plasmids with similar or dissimilar resistance profiles) without first conducting further controlled experiments. Therefore, while our experimental system and model helps to understand the structure of natural host-plasmid networks (Risely et al., 2024) and suggest that networks are formed by forces such as the generality of beneficial plasmids and competition between similar plasmids, further work on simple and complex communities is needed. Furthermore, the extent to which plasmids sharing the same antibiotic resistance genes will affect bacteria-plasmid network structure will depend on the extent of resistance overlap in nature. High quality data addressing this question should be forthcoming as sequencing and bioinformatic tools continue to improve.

Here we have shown that the positive relationship between the fitness effects plasmids have on their hosts and the connectance/generality of of bacteria/plasmid networks is weakened in the face of multiple plasmids conferring similar benefits. Thus, the key to the spread of a plasmid is just the absolute fitness effect it has on a host strain, but that effect relative to other plasmids in the community.. Common antibiotics will select for plasmids that carry similar resistance genes. Thus, in natural communities, overlap in the resistance profiles of plasmids will place an upper bound on the increase in generality resulting from carrying beneficial genes. It is possible that different experimental results would hold where the plasmids come from different Inc groups and/or only a subset are beneficial to hosts. These scenarios present interesting avenues for future research.

## Methods

### Experimental methods

We used experimental communities containing three soil bacterial species with no resistances to the antibiotics Tetracycline, Erythromycin and Gentamicin. The isolates *Bordetella sp*., *Ochrobactrum sp*. and *Pseudomonas sp*. show distinct colony morphologies when plated on KB agar and therefore allowed us to monitor community dynamics. To investigate how the level of plasmid costs or benefits for their bacterial hosts impact bacteria-plasmid network structure, we co-cultured the 3 strains with 3 Inc-P plasmids Pb5, Pb12 (Dröge et al., 2000) and pkjk5::gfp (Klümper et al., 2015). All three plasmids confer resistance to the antibiotic tetracycline. All plasmids can infect all three hosts and because they belong to the same Inc group, co-infections are not possible. The plasmids were chosen because plasmid carriage for each host-plasmid pair can be identified by distinct resistance profiles for PB5, PB12 have and green fluoresces for pkjk5 (see replica plating below).

The presence of tetracycline changes the type of interaction between bacteria and plasmids: In the presence of tetracycline the plasmids are beneficial to their hosts (see fitness effects of plasmids in Figure 1). Since the plasmids also depend on bacteria for replication, under tetracycline exposure this is a mutualistic interaction and beneficial for both.

To set up the host-plasmid communities, we initiated overnight cultures for every host and host-plasmid combinations from freezer stocks. Bacteria with plasmids were cultured on in KB broth with 1ul/ml tetracycline to ensure 100 % plasmid infection rate. Bacteria without plasmids were grown in KB broth. After 12 hours, all overnight cultures were washed 3 times in 1/64 TSB to remove any tetracycline. All cultures were diluted to the same cell density measures by OD600 and mixed as communities to have equal ratios of each host. Therefore, we had two starting communities with the same cell density: one with plasmids and one without plasmids.

Communities with plasmid had overall infection rate at 10 % (3.34 % per plasmid).

We established 6 treatments: no tetracycline (No Tet), 0.1μg/ml tetracycline (Low Tet), 1 μg/ml tetracycline (High Tet), with all three plasmids present/absent. Our experimental microcosms were glass vials with 20 ml 1/64 concentration tryptic soy broth (TSB) with glass beads of radius 1mm in (filled to the top of the fluid content) incubated at 28 degrees statically. Glass beads were used to simulate a structural component such as sand that allow surface growth and increase species persistence in experimental communities (Newbury et al., 2023). We set the experiment up with 10 replicates per treatment and removed any contaminated replicates from the analyses. Every week, we transferred 200μl into new glass vials with fresh medium and beads with the respective tetracycline concentration for each treatment. The experiment ran for 6 weeks, and we sampled each community in week 1 (3 days after the set up) and in week 6. Samples were stored as glycerol stocks and plated on KB Agar at a 10-4 dilution.

Initially 3.34 % of each host population was infected with each of the plasmids. Therefore, host-plasmid networks were at the start of the experiment fully connected with a connectance of 1. At weeks 1 and 6 colony morphology and replica plating on selective KB agar plates was used to determine the prevalence of each bacterium-plasmid combination and construct host-plasmid interaction networks (Figure 2a). We first identified the colonies belonging to the three bacteria on the plates and then plasmid-host associations were revealed by replica plating on selective plates as pB5 and pB12 have distinct resistance profiles: Gentamicin (10 μg/ml) selects for any bacteria carrying the plasmid Pb5, while Erythromycin (80 μg/ml) selects for bacteria carrying the plasmid pB12. Infections with pkjk5 were detected via expression of plasmid-encoded green fluorescent protein (gfp) for all hosts. Replica plating also confirmed that all plasmids were incompatible – no colonies were detected with more than one plasmid, and that tetracycline resistance genes were not integrated into the bacterial chromosomes at any significant rate – all tetracycline resistant colonies hosted plasmids.

### Networks

We used the plotweb function from the package bipartite (Dormann et al., 2008) in R version

4.4.2 (2024-10-31) (R Core Team, 2014) to plot the host-plasmid networks. In Figure 2 we present average networks per treatment with average relative host abundance and plasmid infection rate. For the analysis we uses data for each single replicate community. To test how treatment impacts the generality of the plasmid, we calculated connectance (as the number of realized links divided by the number of all possible links) and the generality of the plasmid per replicate in weeks 1 and 6. For generality, we used G = 2H, with H being the Shannon diversity of interactions for the plasmid. We then used connectance and plasmid generality as response variables to test for the impact of treatment (concentration of tetracycline) and time (weeks 1 and 6) by including the interaction between the factors in the model. We used a Bayesian linear mixed-effects model to examine the effects of treatment, and week on the two network metrics (connectance and plasmid generality). The model was fitted using the brm() function from the brms package (Bürkner, 2021), which employs Hamiltonian Monte Carlo (HMC) sampling to estimate posterior distributions. The fixed effects included treatment and week as well as their interaction (i.e., week * treatment). The treatment variable was a categorical factor with three levels (no, low, high). We included the interaction between week (week 1 and 6) and treatment to test whether the effect of treatment changes over time. We included random intercepts for replicates to account for the hierarchical structure of the data, where observations from the same replicate are expected to be correlated. Weakly informative priors were applied to the fixed effects and random effects. The model was fit using four Markov chains with 2000 iterations, including 500 warmup iterations per chain. The outcome was modeled using a Gaussian distribution. Model diagnostics indicated convergence, with all R-hat values below 1.1 and adequate posterior predictive checks.

We used another model to test how plasmid type might affect their generality, using plasmid specific generality measure as response variable. For this model with a similar structure as above, the fixed effects included treatment, plasmid ID (pb5, pb12, pkjk5), and week, as well as their interactions (i.e., week * treatment * plasID). plasID was included as a categorical factor.

### Relative abundance

In order to visualise relative abundances with some measure of the uncertainty in our data, the typical relative abundance of each host, plasmid and host-plasmid combination was estimated using Dirichlet-multinomial models (Harrison et al., 2020).

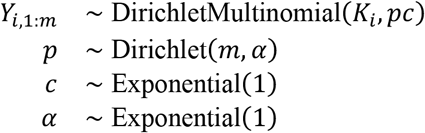

where *Y* is an *n* by *m* data matrix containing rows of count data, *K* is a vector containing the sum of each row of *Y*. The model was implemented in Turing.jl (Ge et al., 2018) using the No U-Turns (NUTS) algorithm (Homan & Gelman, 2014) with 4 chains of 1000 iterations each (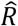 < 1.01 for all parameters).

### Plasmid effects on growth rate

For each host, in each treatment condition (No Tet, Low Tet, High Tet) we estimated the effect of each plasmid by fitting logistic growth curves to optical density (OD) data from growth in 200 μl TSB growth media, at 28°C, with 5 μl of mineral oil suspended on top of the growth media to prevent evaporation. All growth assays were carried out in a single 96-well plate, with 3 replicates per combination of host and plasmid (including no plasmid) in the No Tet and Low Tet conditions, and 2 replicates per host-plasmid combination in the High Tet condition. The reduced number of replicates for the High Tet condition allowed all assays to take place within the same 96-well plate. In the High Tet condition, none of the bacterial strains grew without plasmids, meaning that the benefits of each plasmid were either completely clear on visual inspection, or (in the case of *Pseudomonas sp*. and pb5) nonexistent, as there was still no growth even with the plasmid. The amount of OD data used varied by host species, depending on their growth dynamics – 36, 24 and 96 hours for *Bordetella sp. Ochrobactrum sp*. and *Pseudomonas sp*. respectively. The mean OD for each host-plasmid combination was contrasted with the no-plasmid mean OD values for each treatment with a Bayesian linear model with priors N(0,10) for the intercept (i.e., the no-plasmid mean OD) and the change in mean OD caused by each plasmid and exponential(1) priors for the residual variance. The posterior was inferred via Markov chain Monte Carlo, using the No U-Turns (NUTS) algorithm implemented in Turing.jl in the Julia programming language (Bezanson et al., 2017), with 4 chains of 1000 iterations each (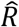 < 1.01 for all parameters). It is possible that the plasmids affected the relationship between population density and OD, though this would not change the interpretation that tetracycline causes the plasmids to be increasingly beneficial. Results are shown in Figure 1.

### Mathematical model

We utilised a model described in detail in (Risely et al., 2024), with the dynamics of bacterial host and plasmid populations given by the following ordinary differential equations describing the change in resources, plasmid-hosting bacterial cells and bacteria without plasmids respectively:

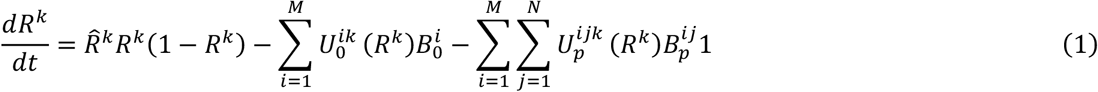

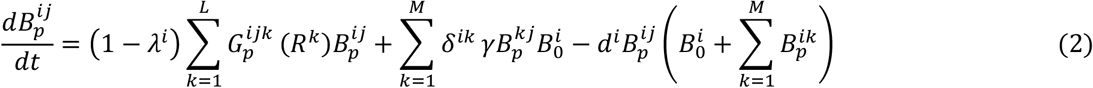

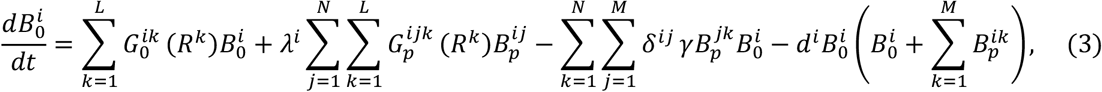

where *R* is the kth resource and 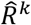 its associated rate of influx, 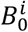 is the density of cells the ith bacterial strain without any plasmid and 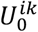 is the function determining the rate at which it consumes the kth resource, 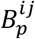 is the density of the ith strain with the jth plasmid and 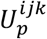 it’s consumption function for the kth resource, *d*^*i*^ is the ith strain’s death rate, *λ*^*i*^ is the ith strain’s rate of loss of plasmids due to segregation, *γ* is the conjugation rate (shared by all hosts/plasmids), *δ* is a square matrix in which all off diagonal elements equal 1 and all diagonal elements equal 10 resulting in 10-fold higher intra-strain than between-strain conjugation rate and 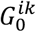 is the function governing the conversion of the kth resource into cells of the *i*^th^ species (plasmid free) and 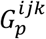 is the same for the *i*^th^ species with the *j*^th^ plasmid.

The model was solved numerically using the fifth-order explicit Runge–Kutta method (Tsitouras, 2011) implemented in the Julia package DifferentialEquations.jl (Rackauckas & Nie, 2017). All model parameters were assigned as described in (Risely et al., 2024)(Risely et al., 2024)(Risely et al., 2024), with the exception that we used only 3 hosts (*M* = 3) to match our experimental conditions. Fitness effects *w* of the plasmids were initially constrained to be negative (as expected in nature (San Millan & MacLean, 2017)(San Millan & MacLean, 2017)(San Millan & MacLean, 2017)) by sampling from a normal distribution Normal(0.985,0.007) truncated between 0 and 1. Then in order to contrast these results with those involving beneficial plasmids we calculated the difference between the lowest *w* value and 1. We added a small number (0.007) to this value, so that its addition to any *w* value would make it positive. Then we added this value either to all *w* values (i.e., all plasmids become beneficial) or to all *w* values related to a single plasmid (i.e., a single plasmid becomes beneficial to all hosts). The former mirroring the experimental set-up with 3 incompatible plasmids all conferring a fitness benefit, and the latter contrasting this with the case where only a single plasmid within an Inc-group confers a significant benefit. There were 864 simulation runs for each type of simulation: all negative, single positive; all positive.

